# Classic articles in Apoptotic Research: A Bibliometric Analysis

**DOI:** 10.1101/2020.08.06.239327

**Authors:** Amadou W. Jallow, Shin-Da Lee, Yuh-Shan Ho

## Abstract

**Background:** Classic articles are defined as research papers with a total citation of one thousand or more. The present study is to identify and analyse the characteristics of the classic articles in apoptosis research.

**Method:** Classic articles with total of 1,000 or more citations from Web of Science Core Collection since year of publication to the end of 2017 were basically assessed regarding their document types, languages, journals, and Web of Science categories within 1900 to 2017.

**Result:** The study showed 418 classic documents in apoptotic research including 260 articles published between 1972 and 2012. The most productive Web of Science category was multidisciplinary sciences. *Nature* published most of these classic articles followed by *Cell*, and *Science*. The most productive country and institution were United States and Harvard University respectively. The author S.J. Korsmeyer from Harvard University was the most productive in apoptosis field and published 13 classic apoptosis articles while the author J.C Reed had more potential to publish classic apoptosis articles in future. The author J.C. Reed and V.A. Fadok had equal potential to publish the same number of classic articles as first- and corresponding-author. Article of Kerr et al. in 1972 was the most popular and cited apoptosis article. The most impact article in 2017 was article entitled “Tumor-associated B7-H1 promotes T-cell apoptosis: A potential mechanism of immune evasion” by Dong et al. in 2002.

**Conclusion:** This study seems to identify the most industrious authors, institutions, and countries in the field of apoptosis research. It also tends to reveal the historical and discoveries related to the pathophysiology of apoptosis as well as the most impact publications on apoptosis studies.

## 1. INTRODUCTION

Decades, after the discovery of the term apoptosis by Kerr, Wyllie, and Currie in 1972 (1) and the involvement of program cell death in the pathogenesis of various diseases and abnormalities in human, gave an extensive interest in the research of apoptosis. John Foxton Ross Kerr was the first to describe the phenomenon of apoptosis and placing the roles of cell death in normal adult mammals and diseases into scientific focus (2). Million cells die in our body every second committing suicide by the mechanism of apoptosis. Apoptosis is also significant for the survival of the body and play vital roles in various developmental process and the immune system (3). Conditions such as cancers, Autoimmune diseases, inflammatory diseases and viral infection can be as a result of apoptosis hindrances and also been known that accumulation of cells is due to decrease cell death (4). Kerr and his group studied the process of cell death in different conditions such as in neoplasms, liver and adrenal injury, and ontogenesis. They found out that the morphological and ultra-structural features displayed by the phenomenon of cell death in each case were similar (5). The occurrence of cell death is interesting where it is being controlled in an event such as embryonic development, metamorphosis and hormone-dependent atrophy but where it is as a result of toxin agents, will undergo different morphology and necrosis will be observed (6). Death of several type of normal and neoplastic lymphoid cell is caused by physiological concentration and glucocorticoid hormones but the mechanisms are unknown (7). A classic article with at least 1,000 citations (8) by Evan in 1992 entitled “Induction of apoptosis in fibroblasts by c-myc protein” was referred as ‘hot paper’ in Scientist (9) because it partly explained the phenomenon of cancer as well as the growth and suicidal pathway of cells. In another classic article, reported that engaging CD3/T-cell receptor complex with anti-CD3 antibodies can induced apoptosis by DNA degradation (10). For many years, neither apoptosis nor programmed cell death was a highly cited term. But after the discovery of the components of cell death, mechanisms and involvement of abnormal cell death in diseases, increased apoptosis researches substantially in the early 1990s (3). Apoptosis may play role in cell turnover in many healthy adult tissues and is accountable for expelling of cells during normal embryonic development (1). In dying epidermal cells, the morphological changes that occur are similar with those in programmed cell death then has essential role in the development of vertebrate embryos and can also contribute in controlling the size of tissues in adult under both normal and pathological state (11). In addition, Kerr and Searle (1973) also presented deletion of cells by apoptosis during castration-induced involution of rat prostate. Bibliometric method can be applied to determine the impact of publications and research groups in their research field (12). It is also an ideal way to quantify the quality of published work for organizations, authors, and countries (13). High quality paper with exemplary for other researchers is considered classic (14). Today, it is essential to determine the countries and institutions research performances using Science Citation Index Expanded or Social Science Citation Index to evaluate their research performances in various aspects (15, 16). In recent years, analysis of classic articles with 1,000 total citations or more in Web of Science categories of surgery (8), psychology (16), neurosciences (17), and orthopedics (18) have been conducted to assess the research performances and productivity among countries and institutions. Publication indicators such as number of total, independent, collaborative, first-author, corresponding-author, and single author classic articles as well as citation indicators including total citations from publication to the end of the most recent year and citations the most recent year only were applied to evaluate publication performances of countries and institutions.

This current study will show the results of classic articles in apoptosis extracted from SCI-EXPANDED. Description and analysis of classic articles in terms of document types, publication language, publication outputs, citation impact and comparison of performances among countries, institutions, and author applying the six publication indicators. We also applied the *Y*-index parameter introduced by Ho (13, 19) to distinguish researchers’ characteristics and who had higher potential in publishing classic articles. Data were clearly drafted and represented in tables and figures for clear evidence and explanation.

## 2. METHODOLOGY

Bibliometric information of research reports was obtained from the online version of Science Citation Index Expanded (SCI-EXPANDED) database in the Web of Science Core Collection of Clarivate Analytics (formerly as Institute for Scientific Information and the Intellectual Property of Thomson Reuters) (updated on February 20, 2019). After our pre-study, keywords such as “apoptosis”, “apoptosissensitive”, “apoptosiss”, “apoptosise”, “apoptosisby”, “apoptosisatase”, “apoptosisinducedby”, “apoptosisrelated”, “apoptosisinduced”, “apoptosisof”, “apoptosisasa”, “apoptosisanducing”, “apoptosisassociated”, “apoptosising”, “apoptosisin”, “apoptosismediated”, “apoptosisl”, “apoptosisis”, “apoptosisinducing”, “apoptotic”, “apoptotically”, “apoptoticMCF-7”, “apoptoticity”, “apoptoticmolecules”, “apoptotickeratinocytes”, “apoptoticinecratic”, “apoptoticassociated”, “apoptoticcells”, “apoptoticphagocytic”, “apoptotical”, “apoptoticrelated”, “apoptotics”, and “apoptoticlike” to be “apoptosis” and “apoptotic” were selected and searched in terms of topic (including title, abstract, author keywords, and *KeyWords Plus*) within the publication years from 1900 to 2017. *KeyWords Plus* is an application for citation indexing using terms extracted from the titles of publications cited by authors in the ISI (now Clarivate Analytics) to be focus on (20, 21). In total, 462,566 documents including 379,030 articles were obtained.

Total citations from Web of Science Core Collection since date of publication to the end of 2017 with the notation *TC*_2017_ (22, 23). Classic articles were defined as *TC*_2017_ ≥ 1,000 (8). Similarly, *C*_2017_ was defined as the total citations within 2017 (13) and was used to characterize the classic articles (8). All the records and citations for each paper was recorded yearly then downloaded and confirmed in spreadsheet software. It was pointed that it is necessary to have a data treatment when using the Web of Science database for bibliometric studies (24).

Thus we manipulated using Microsoft Excel 2016 (25, 26). Publication performance of countries and institutions were analysed by using the six Ho’s publication indicators such as total number of publications (*TP*), independent publications (*IP*), collaborative publications (*CP*), first-author publications (*FP*), corresponding-author publications (*RP*), and single-author publications (*SP*) (12, 27, 28). Affiliations such as England, Scotland, Northern Ireland, and Wales were reclassified to be part of United Kingdom (UK) (27). Affiliations in Fed Rep Ger (Federal Republic of Germany) reclassified to be part Germany (13). The corresponding-author is designated as the “reprint author” in the SCI-EXPANDED database. Therefore, we will hereby be using the term “corresponding-author” in this study (29). In a situation where authorship is unspecified in a single author paper, the author is both the first- and corresponding-author (13). Similarly, as in a single institutional paper, the institution is classified as the first- and corresponding-author institution.

We also applied the *Y*-index indicator proposed by Ho to assess and compare publication potentials of different authors and their contribution characteristic in single index within the same research field. The *Y*-index is related to the number of first-author publications (*FP*) and corresponding-author publications (*RP*). The *Y*-index combines the two parameters *j* and *h*, denoted as (*j*, *h*) (13, 19). This indicator was also being applied to compare authors’ publication performances in classic articles published by American scientists (30).

The *Y*-index (*j*, *h*) is defined as:

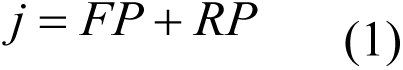

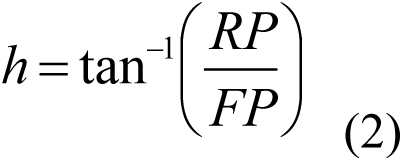

where *FP* is the number of first-author articles; *RP* as number of corresponding-author articles; *j* is the publication potential which is a constant related to publication quantity, and *h* is publication characteristics which can describe the proportion of *RP* to *FP*. The greater the value of *j*, the more the contribution of the first- and corresponding-author articles. Different values of *h* represent different proportions of corresponding-author articles from first-author articles. *h* > 0.7854 indicates more corresponding-author articles; *h* = 0.7854 indicates the same number of first- and corresponding-author articles; and *h* < 0.7854 indicates more first-author articles. When *h* = 0, *j* is the number of first-author articles, and *h* = π/2, *j* is the number of corresponding-author articles.

## 3. RESULTS

### 3.1. Document type and language of publication

We found 418 classic apoptotic publications with *TC*_2017_ ≥ 1,000 within seven document types indexed in the Web of Science. Among the documents types, Article received 260 publications (62% of 418 publications) with a total citation of *TC*_2017_ = 466,231 and review had 151 (36% of 418) publications with *TC*_2017_ = 264,578. The number of citations per publication (*CPP*_2017_ = *TC*_2017_/*TP*) for article and review were 1,793 and 1,752 respectively (Table 1). Document of articles was further studied. All the classic apoptotic articles in SCI-EXPANDED were published in English.

**Table 1.** Characteristics of document types

| Document type | TP | % | AU | APP | $TC_{2017}$ | $CPP_{2017}$ |
| --- | --- | --- | --- | --- | --- | --- |
| Article | 260 | 62 | 2,019 | 7.8 | 466,231 | 1,793 |
| Review | 151 | 36 | 421 | 2.8 | 264,578 | 1,752 |
| Proceedings paper | 4 | 1.0 | 51 | 13 | 7,129 | 1,782 |
| Note | 4 | 1.0 | 34 | 8.5 | 6,172 | 1,543 |
| Editorial material | 2 | 0.48 | 4 | 2.0 | 2,954 | 1,477 |
| Letter | 1 | 0.24 | 6 | 6.0 | 1,172 | 1,172 |
| Book chapter | 1 | 0.24 | 3 | 3.0 | 1,067 | 1,067 |

$TP$ : number of publications; $AU$ : number of authors; $APP$ : number of authors per publication;
$TC_{2017}$ : the total number of citations from Web of Science Core Collection since publication
to the end of 2017; $CPP_{2017}$ : number of citations ( $TC_{2017}$ ) per publication ( $TP$ ).

### 3.2. Characteristics of publication outputs and citation impact

The classic articles were published between 1972 and 2010. The average *TC*_2017_ was 1,793 with maximum of 10,671. The 260 classic articles received a total of 466,231 citations. The distribution of the 260 classic apoptosis articles over years and their *CPP*_2017_ are shown in Fig. 1. The article entitles: “Apoptosis: Basic biological phenomenon with wide-ranging implications in tissue kinetics” (1) published by J.F.R. Kerr, A.H. Wyllie, and A.R. Currie from the Department of Pathology at University of Aberdeen in the UK, was the earliest and only classic article published in 1972. This article shows the highest *TC*_2017_ of 10,671. The year 1995 was the most productive period with 33 articles been published by 49 institutes including Harvard University with seven articles, Washington University (4 articles), and three for each MIT, La Jolla Institute for Allergy and Immunology, and Children’s Hospital in USA respectively. Followed by the year 1998 with 27 articles, 1997 with 26 articles, and 1999 with 21 articles. The article entitled “The landscape of somatic copy-number alteration across human cancers” (31) by 62 authors from USA, Japan, Spain, and Canada with *TC*_2017_ value of 1,450, was the most recent classic article published on apoptosis.

**Figure 1.**
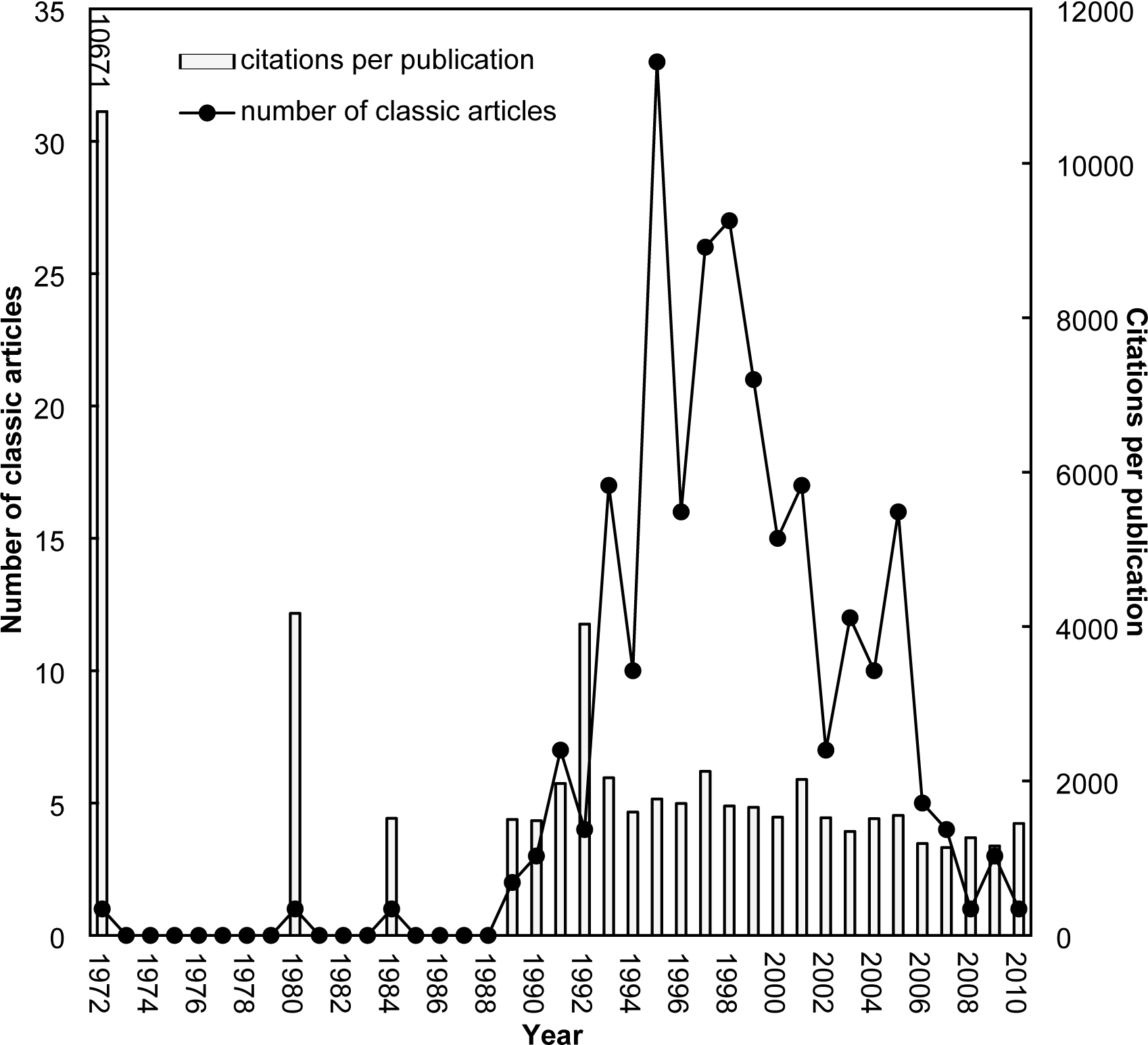
Number of classic articles and citations per publication by year

### 3.3. Journals and Web of Science categories

According to Journal Citation Reports (JCR), it indexes 9,015 journals with citation references across 178 Web of Science categories in the SCI-EXPANDED in 2017. In this context, we found 260 classic apoptosis articles published in 47 journals across 27 Web of Science categories in SCI-EXPANDED. Among the journals, 29 (62% of 47 journals) contained one classic article and six (13%) contained two. A total of 260 articles were published in 47 journals with journal impact factor information in 2017 (*IF*_2017_) and only one article was published in *International Review of Experimental Pathology* which had no journal impact factor after 1996. Figure 2 shows how these *IF*_2017_ are distributed. Table 2 showed that 65% of 260 articles were published in the top four productive journals with *IF*_2017_ higher than 30 (*IF*_2017_ > 30). As listed in the Table 2, *Nature* (*IF*_2017_ = 41.577) published the most classic articles. The *New England Journal of Me*dicine (*IF*_2017_ = 79.260) with two articles had the highest *IF*_2017_ whereas *Journal of Molecular Graphics & Modelling* (*IF*_2017_ = 1.885) had the lowest *IF*_2017_ with only one article.

**Table 2.**
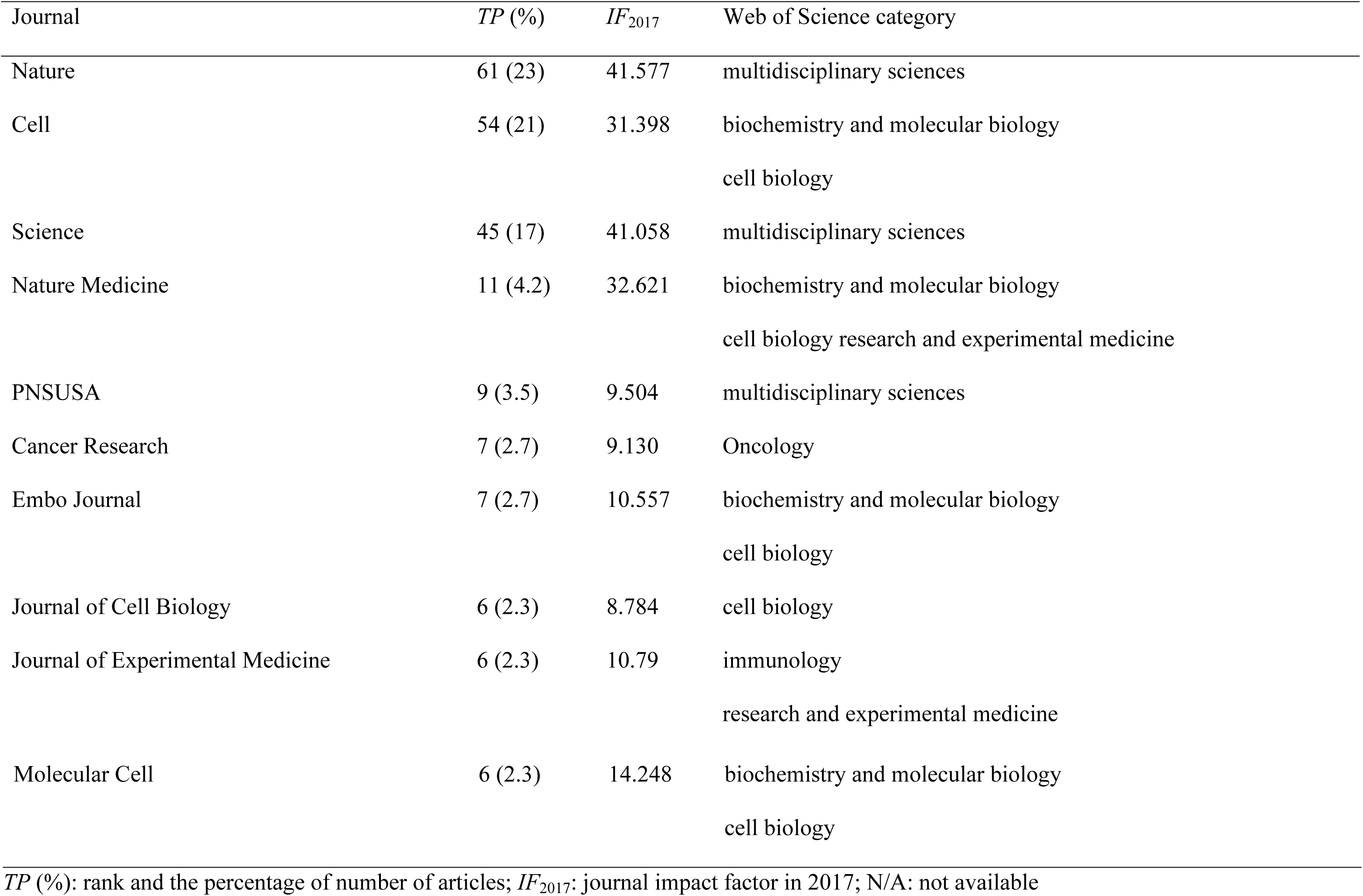
Top 10 most productive journals.

| Journal | *TP (%)* | *IF*<sub>2017</sub> | Web of Science category |
| --- | --- | --- | --- |
| Nature | 61 (23) | 41.577 | multidisciplinary sciences |
| Cell | 54 (21) | 31.398 | biochemistry and molecular biology<br>cell biology |
| Science | 45 (17) | 41.058 | multidisciplinary sciences |
| Nature Medicine | 11 (4.2) | 32.621 | biochemistry and molecular biology<br>cell biology research and experimental medicine |
| PNSUSA | 9 (3.5) | 9.504 | multidisciplinary sciences |
| Cancer Research | 7 (2.7) | 9.130 | Oncology |
| Embo Journal | 7 (2.7) | 10.557 | biochemistry and molecular biology<br>cell biology |
| Journal of Cell Biology | 6 (2.3) | 8.784 | cell biology |
| Journal of Experimental Medicine | 6 (2.3) | 10.79 | immunology<br>research and experimental medicine |
| Molecular Cell | 6 (2.3) | 14.248 | biochemistry and molecular biology<br>cell biology |

*TP* (%): rank and the percentage of number of articles; *IF*<sub>2017</sub>: journal impact factor in 2017; N/A: not available

**Figure 2.**
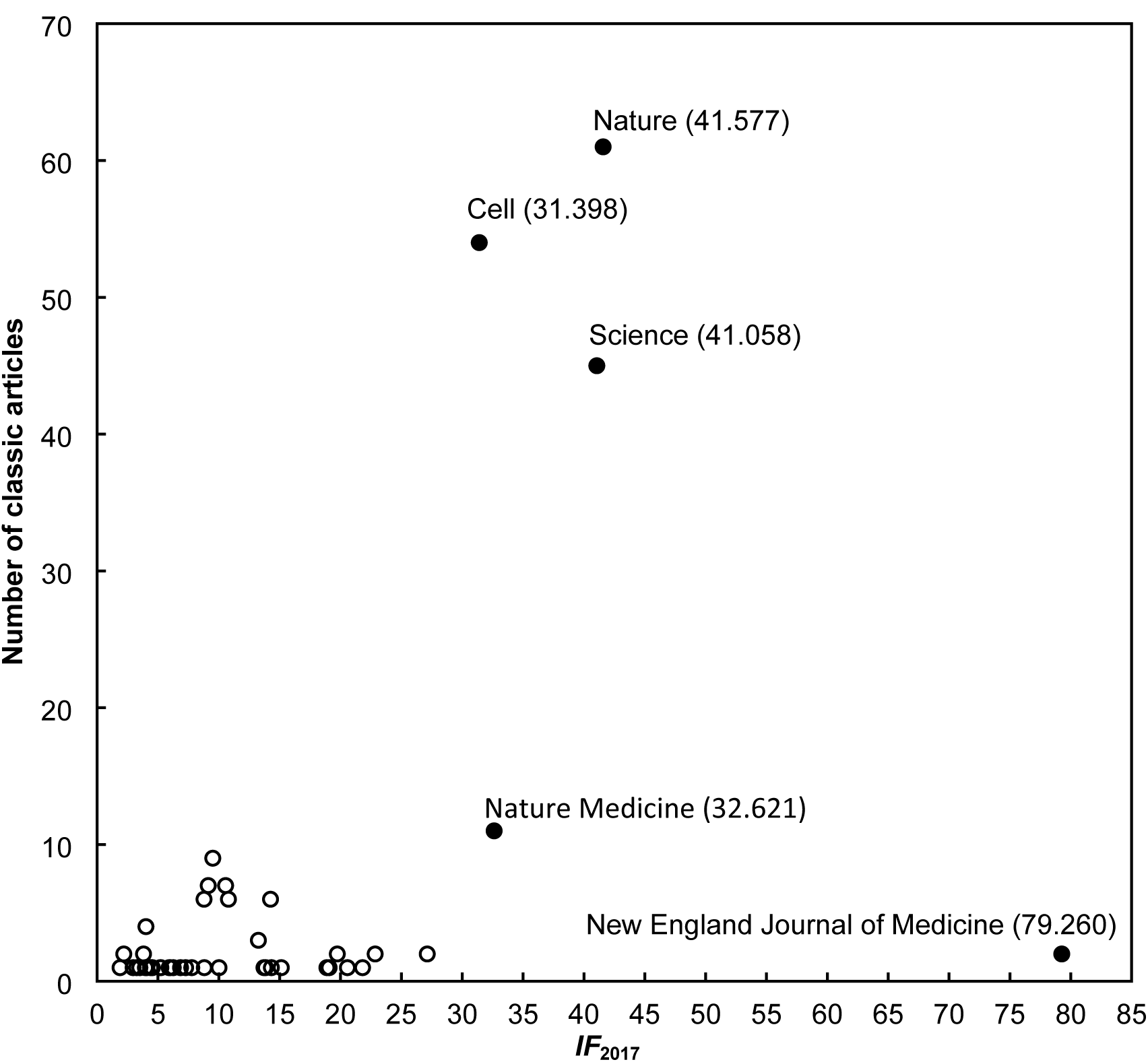
Relation between journal impact factor and number of articles published in journals with *IF*_2017_. *IF*_2017_: journal impact factor in 2017.

In total, 220 articles (85% of 260 articles) were published in the top three categories such as multidisciplinary sciences (44% of 260 articles), cell biology (95 articles; 37%), and biochemistry and molecular biology (93; 36%). Other categories with less articles include, 20 articles in research and experimental medicine and 13 articles in oncology and immunology were found respectively. It has been noticed that journals could be classified in two or more categories in Web of Science, for example *Nature Medicine* was classified in the categories of biochemistry and molecular biology, cell biology, and research and experimental medicine thus the sum of percentages was higher than 100% (19).

### 3.4. Publication Performances: Countries, Institutions, and Authors

The 259 articles with author affiliations from 26 countries, 189 (73% of 259 articles) were single country articles from 13 countries and 70 (27%) were internationally collaborative articles from 25 countries. The top 10 productive countries published seven articles or more with 97% of 259 classic articles as listed in Table 3. The seven major industrialized countries (G7) including the USA, the UK, Japan, Canada, Germany, France, and Italy published majority of the apoptosis papers, contributing 93% of the 259 classic articles, especially the USA with 201 articles (78% of 259 articles). In total, 83 (32% of 259 articles) were single institution articles and 176 (68%) were inter-institutionally collaborative articles including 106 (60% of 176 articles) nationally collaborative articles and 70 (40%) internationally collaborative articles. Table 4 shows the 13 most productive institutions whose authors published no less than nine articles. Harvard University in United State is the most productive institution in the topic of apoptosis. By analysing author, the 260 classic articles of apoptosis were published by 1,687 authors consisting of 239 first-authors and 164 corresponding-authors. Only 194 of the articles had corresponding-author information in SCI-EXPANDED and 66 articles were without corresponding-author information. The author S.J. Korsmeyer from Harvard University published the most 13 classic apoptosis articles with two corresponding-authors and J.C. Reed from Burnham Institute in USA published 12 classic articles including one first-author, five corresponding-author, and one single author articles respectively.

**Table 3.** Top 10 most productive countries

| Country | TP | *TPR* (%) | *IPR* (%) | *CPR* (%) | *FPR* (%) | *RPR* (%) | *SPR* (%) |
| --- | --- | --- | --- | --- | --- | --- | --- |
| USA | 201 | 1 (78) | 1 (76) | 1 (81) | 1 (71) | 1 (70) | 1 (50) |
| UK | 25 | 2 (9.7) | 2 (5.8) | 3 (20) | 2 (6.2) | 2 (7.4) | 1 (50) |
| Japan | 22 | 3 (8.5) | 3 (4.8) | 4 (19) | 3 (5.0) | 3 (5.4) | N/A |
| Canada | 20 | 4 (7.7) | 7 (1.1) | 2 (26) | 8 (1.5) | 6 (2.0) | N/A |
| Germany | 18 | 5 (6.9) | 4 (3.2) | 5 (17) | 4 (3.5) | 4 (3.0) | N/A |
| France | 12 | 6 (4.6) | 11 (0.53) | 6 (16) | 4 (3.5) | 5 (2.5) | N/A |
| Italy | 11 | 7 (4.2) | 7 (1.1) | 7 (13) | 9 (1.2) | 7 (1.5) | N/A |
| Israel | 8 | 8 (3.1) | 5 (2.1) | 9 (5.7) | 6 (1.9) | 7 (1.5) | N/A |
| Sweden | 7 | 9 (2.7) | N/A | 8 (10) | 12 (0.39) | 12 (0.49) | N/A |
| Switzerland | 7 | 9 (2.7) | 5 (2.1) | 12 (4.3) | 6 (1.9) | 7 (1.5) | N/A |

*TP*: number of articles; *TPR* (%): rank and percentage of total number of articles; *IPR* (%), rank and the percentage of single country articles; *CPR* (%), rank and the percentage of internationally collaborative articles; *FPR* (%), rank and the percentage of first-author articles; *RPR* (%), rank and the percentage of the corresponding-authored articles; *SPR* (%), rank and the percentage of single author articles; N/A: not available.

**Table 4.** Top 13 most productive institutions

| Institute | TP | *TPR* (%) | *IPR* (%) | *NPR* (%) | *IPR* (%) | *FPR* (%) | *RPR* (%) | *SPR* (%) |
| --- | --- | --- | --- | --- | --- | --- | --- | --- |
| Harvard University, USA | 36 | 1 (14) | 2 (7.2) | 1 (19) | 1 (14) | 1 (5.4) | 1 (6.4) | N/A |
| Washington University, USA | 15 | 2 (5.8) | 1 (8.4) | 4 (6.6) | 47 (1.4) | 2 (4.2) | 2 (3.9) | N/A |
| Children's Hospital, USA | 14 | 3 (5.4) | N/A | 3 (7.5) | 2 (8.6) | 3 (3.1) | 8 (2.0) | N/A |
| Massachusetts Institute of Technology (MIT), USA | 14 | 3 (5.4) | 16 (1.2) | 2 (10) | 19 (2.9) | 6 (2.3) | 10 (1.5) | 1 (25) |
| Burnham Institute, USA | 10 | 5 (3.9) | 3 (6.0) | 17 (2.8) | 19 (2.9) | 4 (2.7) | 3 (3.4) | 1 (25) |
| Massachusetts General Hospital, USA | 10 | 5 (3.9) | N/A | 4 (6.6) | 11 (4.3) | 16 (1.2) | 10 (1.5) | N/A |
| University of Texas, USA | 10 | 5 (3.9) | 7 (3.6) | 7 (4.7) | 19 (2.9) | 4 (2.7) | 10 (1.5) | N/A |
| Yale University, USA | 10 | 5 (3.9) | 16 (1.2) | 7 (4.7) | 5 (5.7) | 13 (1.5) | 10 (1.5) | N/A |
| Johns Hopkins University, USA | 9 | 9 (3.5) | 5 (4.8) | 28 (1.9) | 11 (4.3) | 6 (2.3) | 6 (2.5) | N/A |
| La Jolla Institute for Allergy and Immunology, USA | 9 | 9 (3.5) | 16 (1.2) | 17 (2.8) | 3 (7.1) | 6 (2.3) | 4 (3.0) | N/A |
| National Cancer Institute, USA | 9 | 9 (3.5) | 16 (1.2) | 11 (3.8) | 5 (5.7) | 49 (0.39) | 40 (0.49) | N/A |
| University of Michigan, USA | 9 | 9 (3.5) | 8 (2.4) | 17 (2.8) | 5 (5.7) | 6 (2.3) | 8 (2.0) | N/A |
| University of Pennsylvania, USA | 9 | 9 (3.5) | N/A | 11 (3.8) | 3 (7.1) | 49 (0.39) | 40 (0.49) | N/A |

*TP*: total articles; *TPR* (%): total number of articles and the percentage of total articles; *IPR* (%): rank and percentage of single institute articles; *NCR* (%): rank and percentage of nationally collaborative articles; *ICPR* (%): rank and percentage of internationally collaborative articles, *FPR* (%), rank and the percentage of first-author articles, *RPR* (%), rank and the percentage of the corresponding-authored articles; *SPR* (%): rank and single author articles; N/A: not available.

The study also went further to focus on the 194 articles (75% of 260 classic apoptosis articles) that had information on both the first- and corresponding-authors in the SCI-EXPANDED applying *Y*-index method. There were 1,405 authors who contributed to the 194 classic apoptosis articles which including information on both the first- and corresponding-authors in the SCI-EXPANDED. Among the authors, 76 authors (5.4% of 1,405 authors) published both first- and corresponding-author articles. In addition, 1,133 authors (81%) had neither first- nor corresponding-author articles.

Figure 5 (*j* Cos *h* and *j* Sin *h* are chosen as the *x* and *y* coordinate axes) displays the distribution of the *Y*-index for the top 272 classic authors who published at least one first-author or corresponding-author articles. Each dot represents one value that could be one author or many authors. The author who had the most potential to publish classic articles in apoptosis field was J.C. Reed (6, 1.373) and V.A. Fadok (6, 0.7854) with the same *j* of 6. Publication characteristics constant, *h*, could help to obtain the different proportion of corresponding-author articles to first-author articles. The advantage of the *Y*-index is that, when *j* of authors is the same, publication characteristics of authors can be indicated by *h* (13, 19). The *j* of, J.Y. Yuan, X.D. Wang, S.W. Lowe, S.J. Martin, H. Ichijo, and T. Takahashi were all the same of 3 (Fig. 5). However *h* of Yuan and Wang was π/2; Lowe, Martin, and Ichijo was 1.107; and Takahashi was 0.4636. Yuan and Wang had only three corresponding-author articles; Lowe, Martin, and Ichijo had greater proportion of corresponding-author articles to first-author articles than Takahashi. Within these 272 classic authors, Takahashi was the only author with *h* < 0.7854 (*h* = 0.4636). Furthermore, A. Brunet and T. Nakagawa with *Y*-index (2, 0) published only first-author articles.

In Fig. 5, the author C. Reed and V.A. Fadok have more potential to publish classic apoptosis article in the future. Takahashi had a low publication characteristic (*h*) and with more first-author articles than corresponding-author articles, indicating that the top productive authors contributing to apoptosis research field were more likely to be designated as the corresponding-authors. The same result was also reported in the classic articles published by American scientists (30) and the highly cited articles in health care sciences and services field (32)

### 3.5. Highly cited articles and high impact articles in the most recent year

The top 10 classic articles with *TC*_2017_ of 4,112 or more were listed in Table 5. Four of the 10 articles were published in *Cell* with *IF*_2017_ of 31.398. One was published in six journals respectively. Article entitled “A rapid and simple method for measuring thymocyte apoptosis by propidium iodide staining and flow-cytometry” (33) published in *Journal of Immunological Methods* with low *IF*_2017_ of 2.190. All the top 10 articles were published by single country in which USA published six, UK published two articles, Israel and Italy each published one article. Out of the 10 top classic articles, three were published by each of the institutions Harvard University and Children’s Hospital in USA. In addition, Children’s Hospital also published the three first-author of the top 10 classic articles. In the field of apoptosis research, the article published by Kerr, Wyllie, and Currie in 1972 was the most frequently cited article with *TC*_2017_ of 10,671 and also the only article that had been cited more than 10,000 times (*TC*_2017_ > 10,000). The only two classic articles categorised in both the top 10 of *TC*_2017_ and *C*_2017_ were “Apoptosis: Basic biological phenomenon with wide-ranging implications in tissue kinetics” (1) by three authors from the UK with *TC*_2017_ = 10,671 (ranked 1st) and *C*_2017_ = 281 (ranked 4th) and “Significance analysis of microarrays applied to the ionizing radiation response” (34) by three authors from USA with *TC*_2017_ = 8,224 (ranked 3rd) and *C*_2017_ = 317 (ranked 2nd).

**Table 5.** Top 10 classic articles with *TC*_2017_ > 4,000

| Rank<br>($TC_{2017}$) | Rank<br>($C_{2017}$) | Article title | Reference |
| --- | --- | --- | --- |
| 1 (10,671) | 4 (281) | Apoptosis: Basic biological phenomenon with wide-ranging implications in tissue kinetics | Kerr et al. (1972) |
| 2 (8,514) | 44 (112) | Identification of programmed cell-death Insitu via specific labeling of nuclear-DNA fragmentation | Gavrieli et al. (1992) |
| 3 (8,224) | 2 (317) | Significance analysis of microarrays applied to the ionizing radiation response | Tusher et al. (2001) |
| 4 (5,359) | 15 (175) | Cytochrome c and dATP-dependent formation of Apaf-1/caspase-9 complex initiates an apoptotic protease cascade | Li et al. (1997) |
| 5 (4,736) | 46 (110) | Bcl-2 heterodimerizes in vivo with a conserved homolog, bax, that accelerates programmed cell death | Oltvai et al. (1993) |
| 6 (4,613) | 90 (70) | Opposing effects of erk and JNK-p38 map kinases on apoptosis | Xia et al. (1995) |
| 7 (4,349) | 38 (120) | Akt phosphorylation of BAD couples survival signals to the cell-intrinsic death machinery | Datta et al. (1997) |
| 8 (4,308) | 11 (181) | Akt promotes cell survival by phosphorylating and inhibiting a forkhead transcription factor | Brunet et al. (1999) |
| 9 (4,171) | 123 (47) | Glucocorticoid-induced thymocyte apoptosis is associated with endogenous endonuclease activation | Wyllie (1980) |
| 10 (4,112) | 68 (86) | A rapid and simple method for measuring thymocyte apoptosis by propidium iodide staining and flow-cytometry | Nicoletti et al. (1991) |

$TC_{2017}$ : the total number of citations from Web of Science Core Collection since publication to the end of 2017; $C_{2017}$ : the total number of citations in 2017 only.

Figures 3 and 4 show the citation historical trends of the top 10 most frequently cited articles (*TC*_2017_ ≥ 4,112) and the top 10 high impact articles in 2017 with *C*_2017_ ≥ 183. Earlier publications, for example Wyllie (1980) (6) and Xia et al.(1995) (35) had a long impact history, but much less impact in 2017 (Fig. 3). Sharply increasing trends of citations after publication year and then reached a citation peak could be found most in the top 10 articles but Kerr et al. (1972) and Wyllie (198) had low citations after publication for more than a decade (Fig. 3). One of the top 10 most impact articles in 2017, was published in the 1970s and the 2010s respectively and others were published in the 2000s (Fig. 4). Articles entitled “Tumor-associated B7-H1 promotes T-cell apoptosis: A potential mechanism of immune evasion” by 13 authors from Mayo Clinic in USA was the most impact article in 2017 with *C*_2017_ of 363. We expect this to receive more citation in the future (Fig. 4). For twenty years, article by Kerr et al. (1972) remained receiving citations less than 50 but rapidly reached a high plateau of 664 citations within 10 years whereas article by Trusher et al. (2001) sharply reached a citation plateau within nine years and depreciated. Moreover, article by Dong et al. (2002) (36)was receiving more citations and is likely to elevate higher in the future.

**Figure 3.**
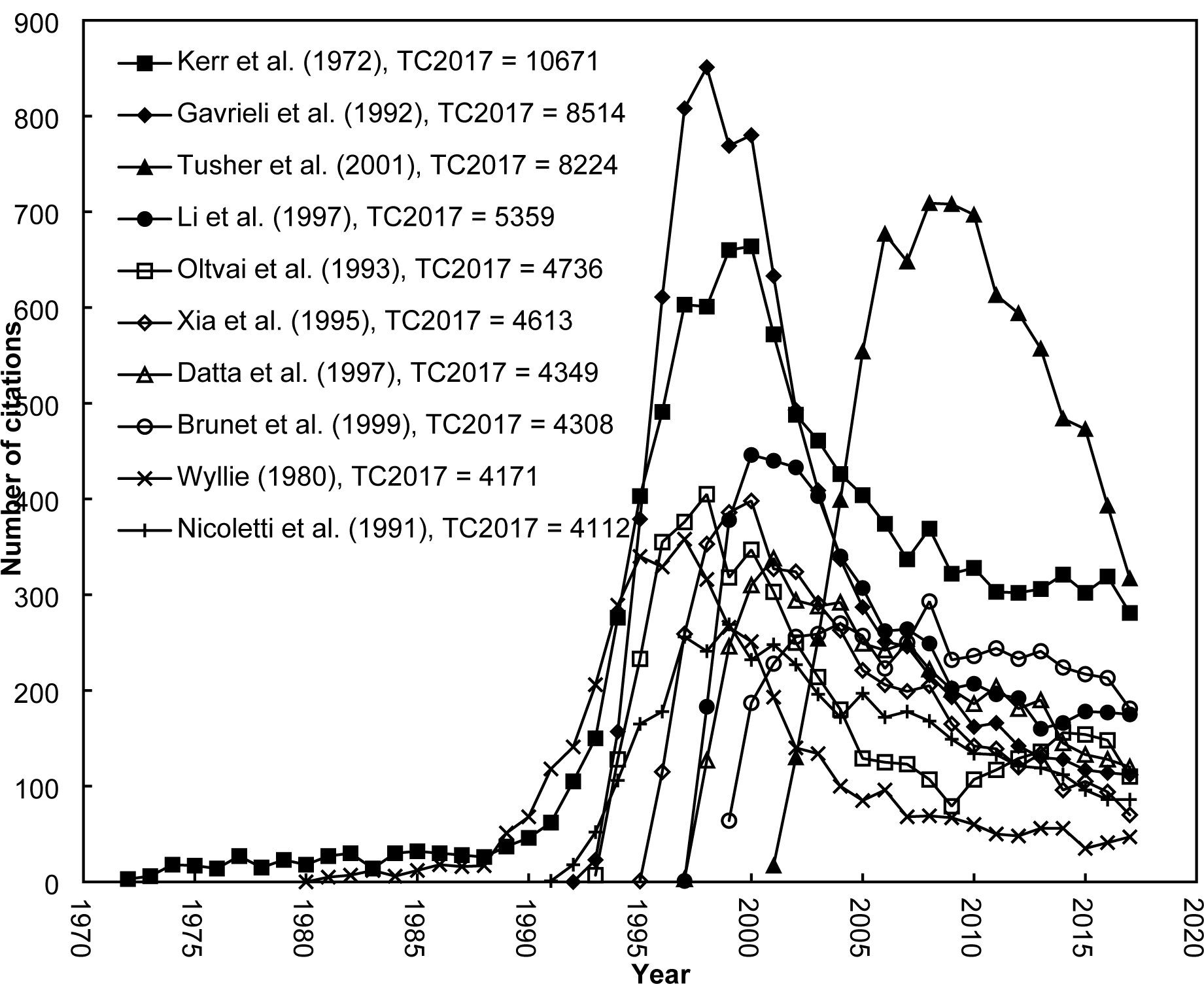
The citation histories of the 10 classic articles with *TC*_2017_ ≥ 4,112. Top 10 classic articles in Apoptotic Research with total citation in 2017 > 4,112. Article list is shown in Table 5.

**Figure 4.**
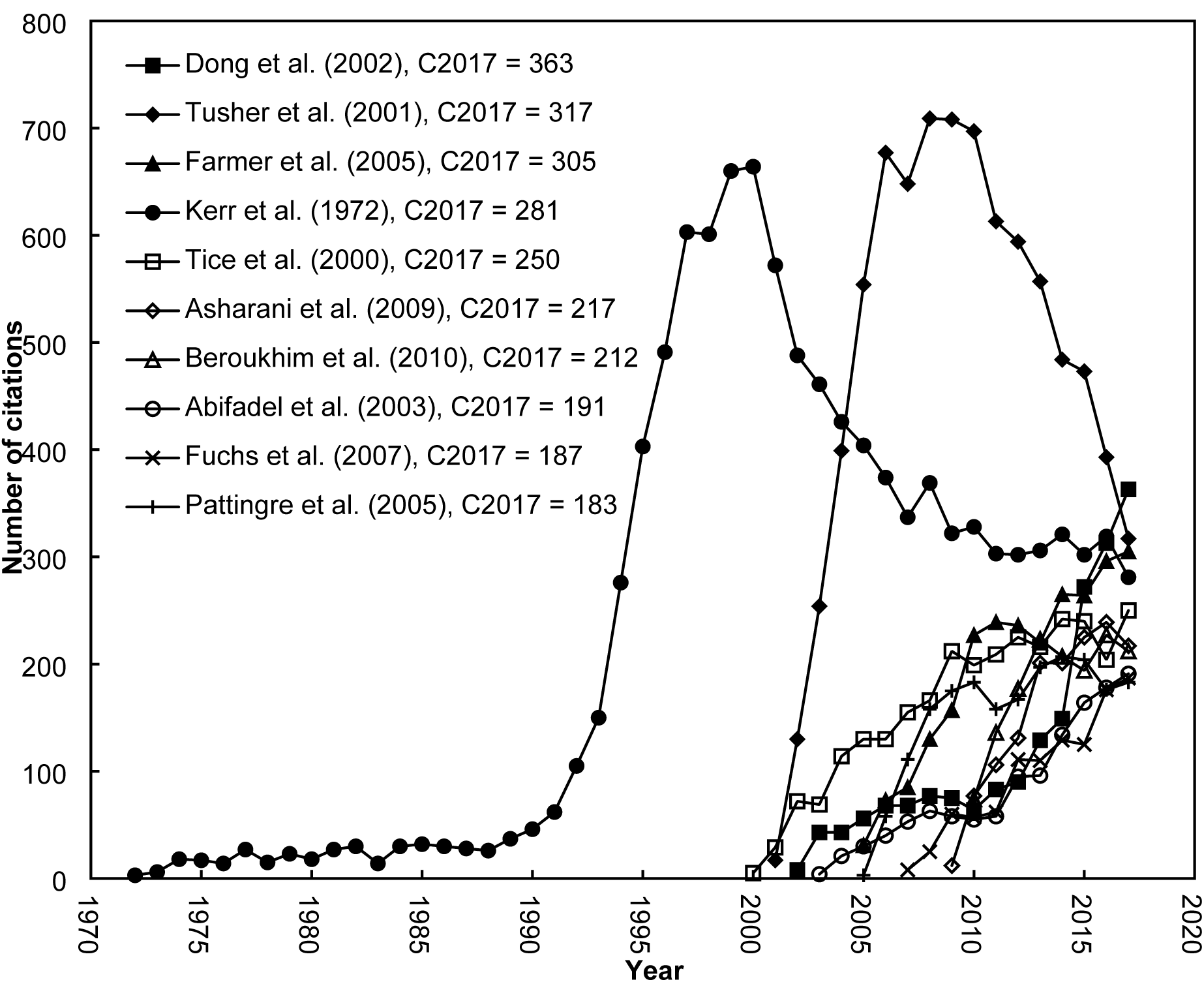
The current trend of the top 10 most cited articles in Apoptotic Research in 2017 ≥ 183. *C*_2017_: the total number of citations in 2017 only.

**Figure 5.**
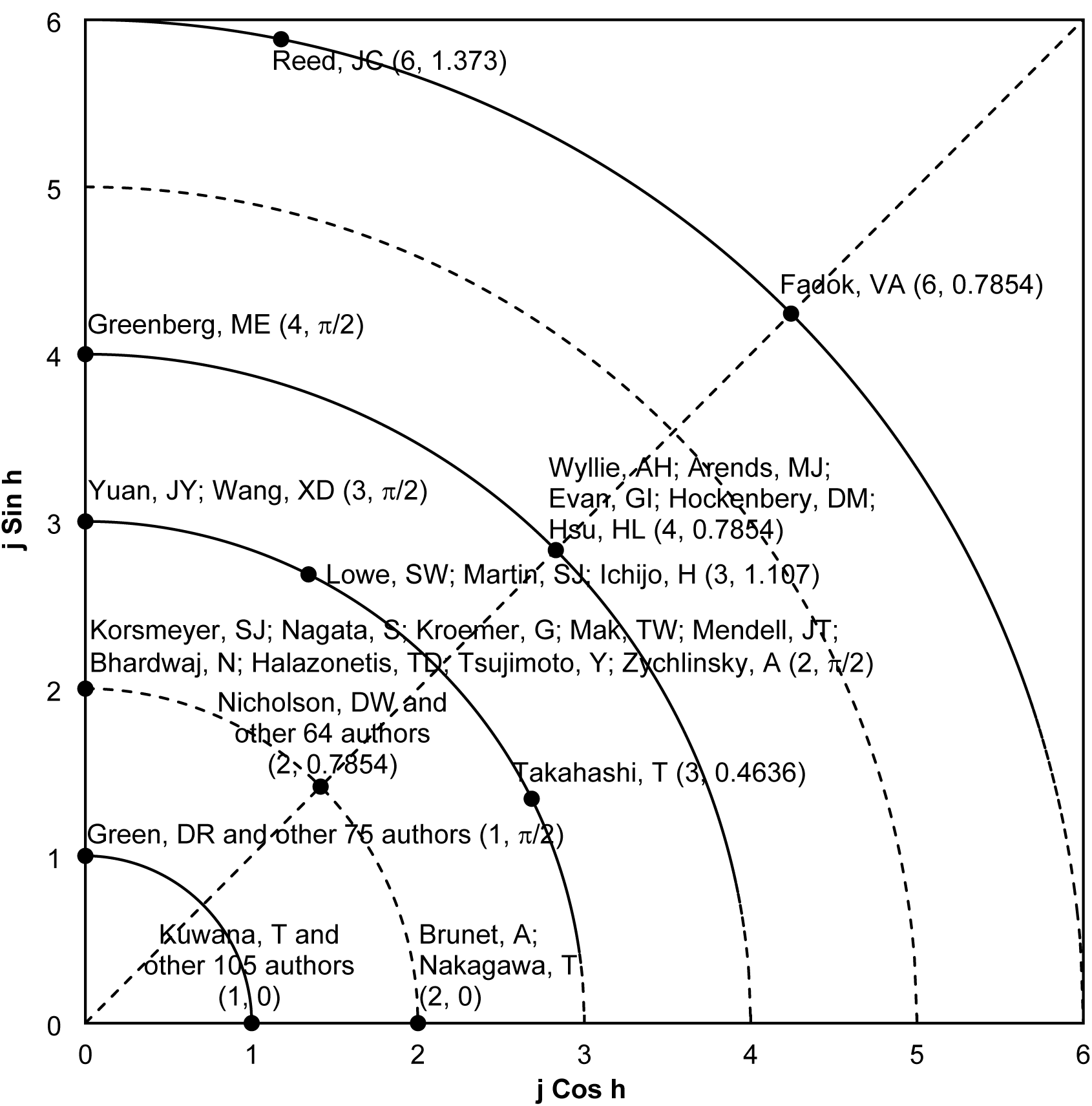
Y-index of the 272 authors in Apoptotic Research. Figure 5 displays the distribution of the Y-index for the top 272 classic authors in Apoptotic Research who published at least one first-author or corresponding-author articles. j Cos h and j Sin h are chosen as the x and y coordinate axes respectively. j represents sum of the first-author articles and corresponding-author articles; h indicates constant characteristic of publication. Each dot represents one value that could be one author or many authors.

## 4. DISCUSSION

### Major findings

There were 418 classic apoptosis publications in SCI-EXPANDED within seven document types. Articles were found slightly higher in citations per publication than that of reviews. All classic articles were published in English. The most classic articles were published in 1995 while article in 1972 had the highest citations per publication. The 260 classic articles were published in 47 journals listed in 27 Web of Science categories. Nature, Cell, and Science were the three most productive journals. 44% of articles were published in Web of Science category of multidisciplinary sciences. The group of seven (G7) countries (the USA, the UK, Japan, Canada, Germany, France, and Italy) published the most articles and ranked in top seven. Thirteen institutions published nine or more classic articles each, and all of these institutions were located in the US. Harvard University ranked top in terms of the number of classic articles published, nationally collaborative, internationally collaborative, first-author, corresponding-author articles while Washington University ranked top in single institute articles. S.J. Korsmeyer published the most classic apoptosis articles while J.C. Reed and V.A. Fadok had the highest potential to publish classic articles in apoptosis field. The article published by Kerr, Wyllie, and Currie in 1972 had the highest number of citations from Web of Science Core Collection since publication to the end of 2017 and the total number of citations in 2017. This could be as a result of their paper describing the nature of apoptosis in embryogenesis, cancer growth, and tissue remodelling during healing or functional regression.

The most frequently cited articles are usually published in journals with high impact factor (*IF*) (37). Nature, Cell, Science and Nature Medicine were the most productive journals.

Theses journals received more classic articles with high IF 2017. Majority of the articles were published in the top three subject categories including multidisciplinary sciences, cell biology, and biochemistry and molecular biology.

In recent year, Ho’s group introduced a relationship between number of classic articles in a year (*TP*) and their citations per publication (*CPP*_year_ = *TC*_year_/ *TP*) by decade as a figure in medical related fields, for example surgery (8), psychology (16), and neurosciences (17).

In terms of country publication performances with the indicators *TP*, *TPR*, *IPR*, *FPR*, and *RPR* introduced by Ho and his group. We found USA as the most industrious followed by UK and then Japan.

Harvard University was considered the most productive and then follow by Washington University with far distance. Children’s Hospital and Massachusetts Institute of Technology (MIT) both ranked equal in terms of total publication (*TP*). Harvard University has the authors name S.J Korsmeyer who published the most 13 classic apoptosis articles with two corresponding-authors. Therefore, we may conclude by recommending Harvard University as the best institution for apoptosis studies worldwide.

The articles in table 5 were the most highly cited with *TC*_2017_ > 4,000. This study has shown that these articles were highly cited to foster more understanding on the pathophysiology of apoptosis. The most highly cited articles will be explained as follows:

1. *Apoptosis: Basic biological phenomenon with wide-ranging implications in tissue kinetics Kerr et al. (1972) (1)* This was the first classic article on apoptosis. The researchers described the nature of apoptosis and its involvement in certain disease conditions. The term apoptosis became popular after this article was published then follow by the discoveries of latest technologies to detect apoptosis in the 1990s.
2. *Identification of programmed cell-death Insitu via specific labeling of nuclear-DNA fragmentation (Gavrieli et al., 1992)* (38): This article was the fastest growing in citations and reached high plateau within a decade. In this article, the researchers developed an observation technique to identify apoptosis (cell death) through staining a fragmented DNA. It was concluded that DNA fragmentation is associated with programmed cell death and therefore contributing to the gate way for more apoptosis studies. This explains why it was rapidly cited.
3. *Significance analysis of microarrays applied to the ionizing radiation response (Tusher et al., 2001)* (34: The Significance analysis of microarrays has paved way for researchers to further investigate genes and their functions. This article introduced a method called Significance Analysis of Microarrays (SAM) to determine genes expression at different biological states. Cells were exposed in ironizing radiation to examine their respond by measuring with microarray. Genes that involves in regulating cell cycle and apoptosis (cell death) were identified. Interestingly, genes that repair DNA damages were also found. This article contributed greatly for apoptosis studies and was swiftly cited to reach a high peak. Furthermore, it was the most cited article in 2017.
4. Cytochrome c and dATP-dependent formation of Apaf-1/caspase-9 complex initiates an apoptotic protease cascade (Li et al., 1997) (39): During the initiation of apoptosis, Cytochrome c is released from the mitochondria to bind with procaspase 9 and Apaf-1 for caspase 9 activation. Activated caspase 9 will trigger caspase 3 to execute apoptosis (cell death). This article was highly cited for describing the mitochondria apoptotic pathway.
5. *Bcl-2 heterodimerizes in vivo with a conserved homolog, bax, that accelerates programmed cell death (Oltvai et al., 1993)* (40): The role of Bcl-2 protein in initiating apoptosis was also investigated in the article. The Bcl-2 interacts with other components for example Bax to progress apoptosis. This article was also highly cited.
6. *Opposing effects of erk and JNK-p38 map kinases on apoptosis (Xia et al., 1995)* (35): This study demonstrated the effect of mitogen-activated protein (MAP) kinase family members such as the extracellular signal-regulated kinase (erk), JNK (c-JUN NH2-terminal protein kinase), and p38, in apoptosis. It was reported that activation of JNK and p38 and simultaneously inhibition of ERK can induce apoptosis. Therefore, a balance between activated ERK and stress-activated JNK-p38 pathways may opposed apoptosis.
7. *Akt phosphorylation of BAD couples survival signals to the cell-intrinsic death machinery (Datta et al., 1997)* (41): Serine-threonine kinase (Akt) plays a very significant role in cell survival but its mechanism is not well known. This article studied the function of akt in apoptosis. Its showed that growth factors can activate akt to phosphorylate with Bad of the Bcl 2 family proteins to inhibit apoptosis and promotes cell survival.
8. *Akt promotes cell survival by phosphorylating and inhibiting a forkhead transcription factor (Brunet et al., 1999)* (42): The purpose of this study on the role of akt in apoptosis was also similar to that of article by Datta et al. (1997). Surprisingly, this article took shorter period to accumulate more than one hundred citations compared to other articles. It could be due to its extensive study on the transcription factor (FKHRL1) related to apoptosis. Suppression of apoptosis transcription factors by Akt will promote cell survival.
9. *Glucocorticoid-induced thymocyte apoptosis is associated with endogenous endonuclease activation (Wyllie, 1980)* (6): This was the second oldest apoptosis article after the first article of Kerr et al. (1972). The author was also part of the group in the first article. This study investigated endogenous endonuclease in apoptosis. Activation of the endonuclease was found to induced chromatin cleavage for DNA degradation (apoptosis). Therefore, inhibiting endogenous endonuclease could be a strategy to prevent apoptosis. This article was highly cited in the 1990s.
10. *A rapid and simple method for measuring thymocyte apoptosis by propidium iodide staining and flow-cytometry (Nicoletti et al., 1991)* (33): Development of new apoptosis detection techniques in the 1990s has also contributed to the increased of apoptosis studies. In this article, the researchers developed a simple technique to calculate the percentage of apoptotic cells with flow cytometry. Those apoptosis detection techniques before 1990 were unable to measure percentage of apoptosis cells in a miscellaneous cell group.

The ten most highly cited articles have revolutionized the pathophysiology of apoptosis and pioneered researchers in apoptosis studies. Above all, apoptosis was described as a programmed phenomenon by Kerr et al., (1972). The article of Oltvai et al. in 1993 (40) and Li et al. (1997) (39) investigated the initiation of apoptosis. They reported that incorporation of members in Bcl-2 protein family group (Bad, Bid, Bax, and Bim) will trigger cytochrome c released to bind with caspase 9 and the adaptor protein Apaf-1 for caspase 3 activation. Activated caspase 3 is the main culprit for apoptosis (cell death). Since apoptosis is a regulatory phenomenon, other studies were focus on the anti-apoptotic regulatory mechanism. Bcl-2 proteins family such as Bcl-2 and Bcl-xL are the anti-apoptotic components and located outside wall of the mitochondria to inhibit cytochrome c released. The article by Datta et al. (1997) (41) and Brunet et al. (1999) (42) described how the apoptotic proteins phosphorylate with other pro survival components including Akt (Protein kinase B) to prevent cell death and promotes cell survival.

Like in Fig. 3, a growth number of apoptosis papers and citations were received between the 1990s and 2000s. Recently, citation trend for all the top ten articles ultimately deteriorated and we believed it could be as a result of research diversity.

In Fig. 4, apoptosis articles published in the 2000s were high impact and article by Dong et al. (2002) (36) was the most impact in 2017. The article of Dong et al. (2002) (36)explained the association of cancel cells with T cell apoptosis for cancer immunotherapy. Due to the increase in cancer studies, we expect the article to receive more citation in the future.

In this study, applying the *Y*-index method was able to help in identifying the most industrious and well-known authors in the research of apoptosis. The memos of few greatest authors who contributed immensely in the field of apoptosis will be discuss as follows.

#### Stanley J. Korsmeyer (1950-2005)

Stanley was a great professor at Harvard medical university with a scientific visionary and driving force for eliminating cancer. Stanley was regarded as the trailblazer in the understanding and treatment of cancer. His contributions and discoveries in apoptosis have revolutionized the understanding of cancer pathophysiology. At his time, he suggested that inducing apoptosis by either activating the pro-apoptotics protein (*Bax*) or blocking the anti-apoptotic genes (*Bcl-2*) may enhances cancer cells for self-destruction (43, 44). In this research, Stanley published most of the classic articles in apoptosis research.

#### John C. Reed (1958, age 61 years)

John left a legacy remaining as unforgettable experience to his colleagues in the laboratory. He is from Burnham Medical Research Institute in united states and a pioneer in apoptosis studies with cancer. In the 1990s, he was regarded as the top biomedical researchers worldwide for his research impact on apoptosis (45). After Stanley, he recorded the most classic apoptosis articles. John dedication and motivation toward his collaborators was adorable and this study has also identified that as he published the most classic corresponding-author articles. In our indicator (*Y*-index), John has more potentials to publish classic apoptosis articles.

#### Scott W. Lowe (1963, age 56 years)

Scott is a biomedical science researcher from Memorial Sloan Kettering Cancer Center and Howard Hughes Medical Institute Investigator. He was interested in studying oncogenes to attack cancer. In his studies on oncogenes, he encountered with a gene called p53 responsible for inhibiting tumour growth. The gene p53 was recommended as the hallmark to onset apoptosis and recently researchers are targeting p53 gene to treat cancer. Scott was recognised for his tremendous effort on cancer and apoptosis (46). In this study, he published the most three first-author articles with K. Kuida.

#### Julian Downward (1960, age 59 years)

He is also another top researcher with scientific visionary in the area of cancer. Julian is from Francis Crick Institute and Senior Group Leader at the Institute of Cancer Research. He was known for his contribution for enlighten cellular signal transduction pathways on oncogenes (47). John potentials was recognised when this study reported him to be among the four authors to publish a single author article.

